# Oxidative stress potentiates the therapeutic action of a mitochondrial complex I inhibitor in MYC-driven B-cell lymphoma

**DOI:** 10.1101/2022.06.21.497021

**Authors:** Giulio Donati, Paola Nicoli, Alessandro Verrecchia, Veronica Vallelonga, Ottavio Croci, Simona Rodighiero, Matteo Audano, Laura Cassina, Aya Ghsein, Giorgio Binelli, Alessandra Boletta, Nico Mitro, Bruno Amati

## Abstract

*MYC* is a key oncogenic driver and an adverse prognostic factor in multiple types of cancer, including diffuse large B-cell lymphoma (DLBCL). Yet, *MYC* activation also endows cancer cells with a series of metabolic dependencies, which can provide strategic points for targeted pharmacological intervention. We recently reported that targeting the mitochondrial electron transport chain (ETC) complex I with the small molecule inhibitor IACS-010759 selectively killed MYC-overexpressing lymphoid cells. Here, we unravel the mechanistic basis for this synthetic-lethal interaction and exploit it to improve the anti-tumoral effects of ETC inhibition. In a mouse B-cell line, MYC hyperactivation and IACS-010759 treatment added up to induce oxidative stress, with consequent depletion of reduced glutathione and lethal disruption of redox homeostasis. This effect could be enhanced by targeted pharmacological intervention, with either inhibitors of NADPH production through the pentose phosphate pathway, or with ascorbate (vitamin C), known to contribute pro-oxidant effects when administered at high doses. In these conditions, ascorbate synergized with IACS-010759 to kill MYC-overexpressing cells *in vitro* and reinforced its therapeutic action against human B-cell lymphoma xenografts. Hence, ETC inhibition and high-dose ascorbate might improve the outcome of patients affected by high-grade lymphomas and other MYC-driven cancers.

**Key point #1:** MYC and the ETC complex I inhibitor IACS-010759 elicit different reactive oxygen species (ROS) that cooperate to disrupt redox homeostasis.

**Key point #2:** Further boosting of oxidative stress with high doses of ascorbate increases the killing of MYC-driven lymphoma xenografts by IACS-010759.

## INTRODUCTION

The *MYC* proto-oncogene and its product, the MYC transcription factor, have a central role in cellular growth control and are widely deregulated in many cancers, where they contribute to many facets of malignant transformation and are generally recognized as adverse prognostic factors^1-3^, as best exemplified by diffuse large B-cell lymphoma (DLBCL)^4-7^. *MYC*-driven tumors also show oncogene addiction, indicating that MYC – and presumably a subset of its target genes – are required for tumor maintenance, based on a diversity of cell-intrinsic and -extrinsic mechanisms^3,7^. Besides multiple attempts to target MYC directly^8,9^, much effort in the field was aimed at the identification of synthetic-lethal interactions as means to develop targeted therapies against MYC-associated tumors^7,10^. Among these, we and others identified pharmacological inhibition of the mitochondrial ribosome with the antibiotic tigecycline as a therapeutically viable strategy against MYC-driven lymphoma^11-13^. Inhibiting mitochondrial protein synthesis interferes with the assembly of Electron

Transport Chain (ETC) complexes and ultimately with Oxidative Phosphorylation (OxPhos)^14^, suggesting that the latter might also be a limiting activity – and hence a direct therapeutic target – downstream of MYC. In line with this concept, we reported that MYC- and OxPhos-associated gene signatures were highly correlated in DLBCL and that a specific inhibitor of ETC complex I, IACS-010759^15^, selectively killed MYC-overexpressing cells by inducing intrinsic apoptosis^16^. In line with this result, the anti-apoptotic protein BCL2 counteracted killing by IACS-010759; reciprocally, the BCL2 inhibitor venetoclax strongly synergized with IACS-010759 against MYC/BCL2 double-hit lymphoma (DHL)^16^, as also reported with tigecycline^13^. Hence MYC sensitized cells to OxPhos inhibition, providing new opportunities for targeted drug combinations against aggressive DLBCL subtypes, and possibly other MYC-associated tumors.

Here, we set out to characterize the mechanistic basis for the synthetic lethal interaction between oncogenic MYC and IACS-010759. We report that MYC hyperactivation and pharmacological inhibition of the ETC coordinately disrupt redox homeostasis, further sensitizing cancer cells to the action of pro-oxidant drugs, with significant improvement of therapeutic efficacies.

## RESULTS

### Disruption of redox homeostasis sensitizes MYC-overexpressing cells to ETC inhibition

We previously showed that ectopic activation of the 4-Hydroxytamoxifen (OHT)-responsive MycER™ chimaera (MycER) in the lymphoid precursor cell line FL5.12 (hereafter FL^MycER^ cells) augmented OxPhos-associated gene expression and respiratory activity, and sensitized cells to apoptotic killing by the ETC complex I inhibitor IACS-010759^16^. To unravel possible MYC-effector pathways underlying this effect, we further queried our previous transcriptomic data^16^. This analysis pointed to the Nrf2-mediated oxidative stress response as the top activated pathway in OHT-treated FL^MycER^ cells, regardless of IACS-010759 (Figure 1A). Consistent with this finding, OHT treatment increased both total and nuclear Nrf2 levels (Figure 1B), alongside a slight induction of the mRNA (1.25-fold), as assessed by RNA-seq^16^. While also inducing ROS^17^ (see below), IACS-010759 had the opposite effect on Nrf2 protein levels.

**Figure 1.**
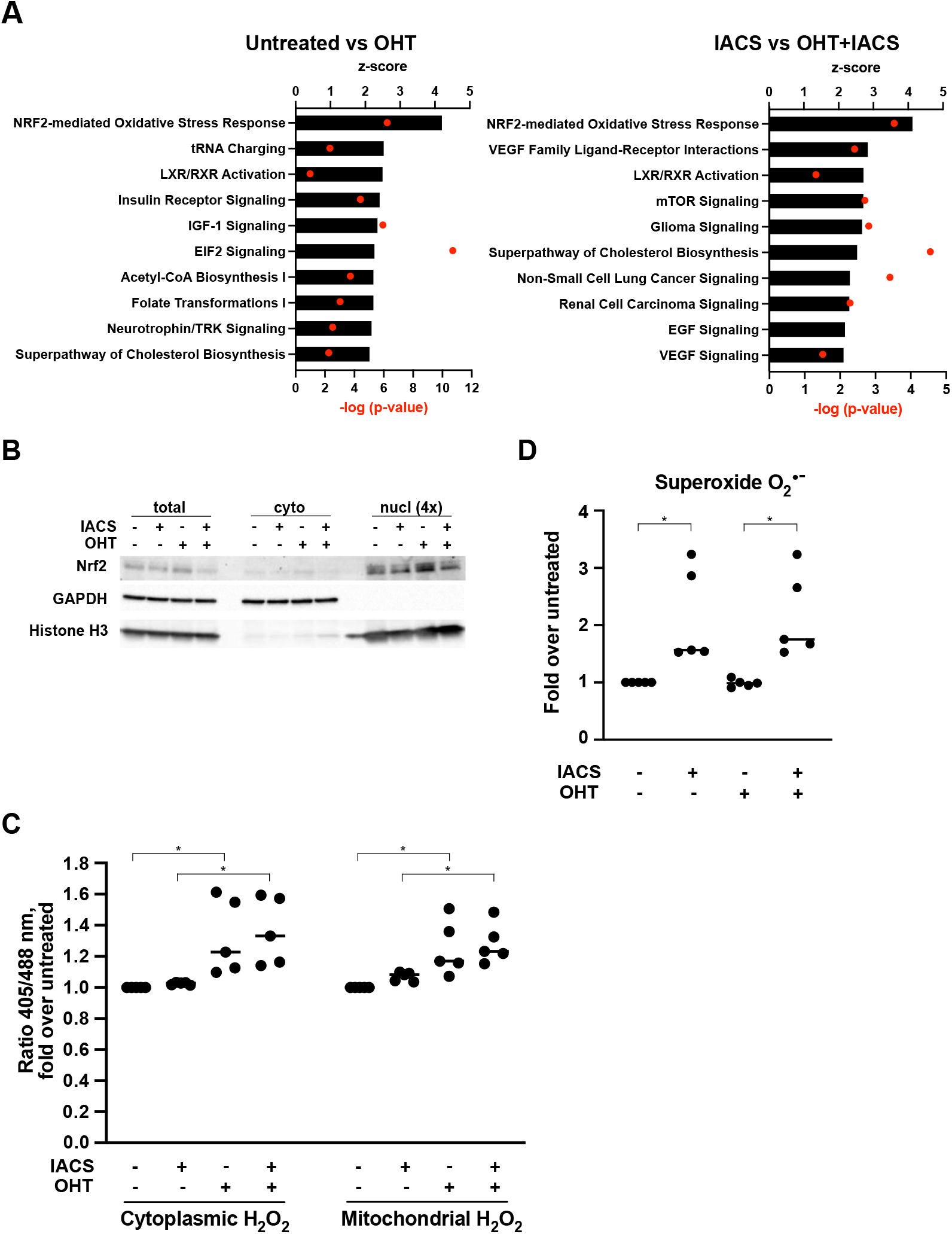
MycER activation causes ROS production and induces an oxidative stress response. FL^MycER^ cells, primed or not with OHT (48h), were treated with 135 nm IACS-010759 (IACS) for 24 (**A**) or 48 hours (**B-D**). (**A**) Top 10 OHT-activated pathways (highest z-score), as determined by IPA Core analysis on DEGs from cells treated or not with OHT (72 hours), either alone (left) or in the presence of IACS-010759 (right). (**B**) Total, cytoplasmic and nuclear Nrf2 levels in FL^MycER^ cells treated as indicated. For the nuclear fraction (right), 4 equivalents of the lysate used for total and cytoplasmic fractions were loaded. (**C**) H2O2 quantification, expressed as fold-increase of the 405/488 nm fluorescence ratio in treated vs. untreated FL^MycER^ cells, expressing either the cytoplasmic (left) or mitochondrial (right) roGFP2-ORP1 biosensor. A representative example of the data is provided in Supplemental Figure 1A. (**D**) Superoxide anion O_2_^•-^ production in treated vs. untreated FL^MycER^ cells, based on dihydroethidium staining. * p ≤ 0.05.

Nrf2 is a transcription factor that under normal growth conditions is bound in the cytoplasm by the Keap1-containing ubiquitin ligase complex and rapidly targeted to degradation. Oxidative stress triggers the release of Nrf2 from Keap1, resulting in its stabilization and migration to the nucleus, where it promotes an antioxidant transcription program^18^. MYC may promote Nrf2 activity at two levels: indirectly through the production of reactive oxygen species (ROS)^19,20^, or through direct transcriptional activation of the *Nrf2* locus^21^. Through the latter, MYC activation may also lead to ROS reduction, as reported in some settings^22^.

Based on the above observations, we sought to address whether – and how – modulation of oxidative stress may underlie the sensitization of MYC-overexpressing cells to IACS-010759. We first addressed the levels of hydrogen peroxide (H_2_O_2_), the most abundant cellular ROS, in FL^MycER^ cells expressing either the cytoplasmic or mitochondrial variants of the H_2_O_2_ biosensor ORP1-roGFP2^23^: as assessed by the 405/488 nm fluorescence ratio, MycER activation led to increased H_2_O_2_ levels in both compartments, while IACS-010759 treatment showed negligible effects (Figure 1C and Supplemental Figure 1A). Most relevant here, H_2_O_2_ is one of the species that can activate Nrf2^18^, and may thus mediate this effect of MycER (Figure 1B). Second, we monitored the superoxide anion O_2_^•-^, produced following impairment of the mitochondrial ETC, and in particular of complex I^24^. Indeed, quantification with the fluorescent probe dihydroethidium^25^ revealed that, contrary to H_2_O_2_, O_2_ ^•-^ was induced upon IACS-010759 treatment, with no significant contribution from MycER (Figure 1D). Notably, while O_2_ ^•-^ is less stable and abundant than H_2_O_2_, the reaction between those two species can produce the highly reactive ROS Hydroxyl radical^26^. Thus, MycER activation and ETC inhibition drive the production of distinct oxidative species, respectively H_2_O_2_ and O_2_^•-^, which may cooperate toward cell killing.

To further monitor the oxidative stress induced by OHT and IACS-010759 in FL^MycER^ cells, we quantified the ratio of reduced to oxidized glutathione (GSH and GSSG, respectively) after 40 hours of IACS-010759 treatment, a time preceding overt cell death (observed at ca. 48h)^16^. Remarkably, each agent alone caused a moderate decrease in GSH/GSSG ratio, which became more pronounced in double treated cells (Figure 2A). Concomitant with these changes, the net levels of both GSH and GSSG were increased in treated cells (Figure 2B), most likely reflecting the activation of compensatory mechanisms to maintain redox homeostasis and favor survival.

**Figure 2.**
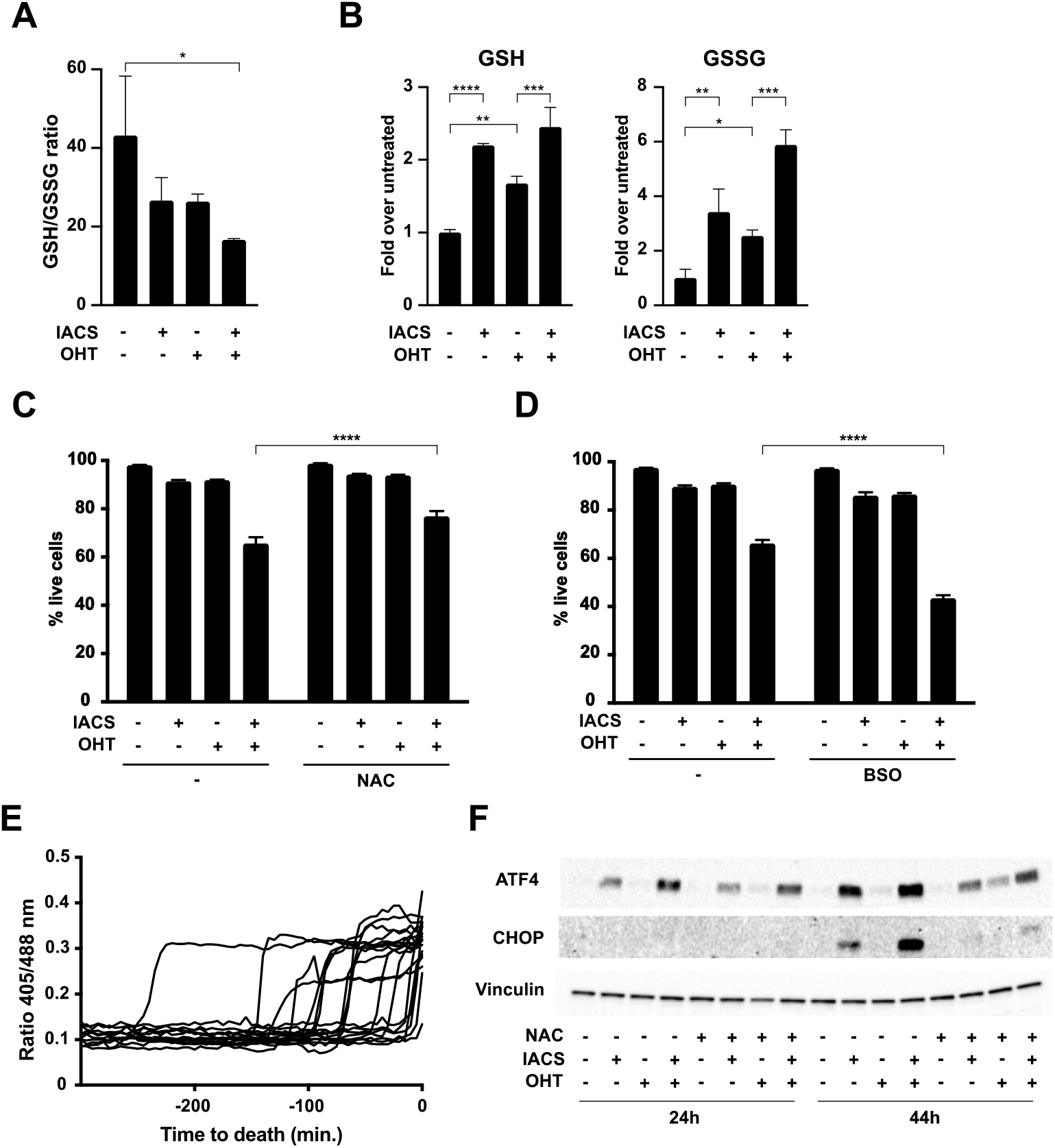
Disruption of redox homeostasis by IACS-010759 induces cell death in Myc-overexpressing cells. FL^MycER^ cells were primed with 100 nM OHT (48h) and/or treated with 135 nM IACS-010759, as indicated. (**A**) Glutathione redox state, given as reduced to oxidized ratio and glutathione quantification in FL^MycER^ cells, measured after 40 hours of IACS-010759 treatment. Cell viability, as determined by PI staining after 48 hours of IACS-010759 treatment, with or without addition of 10 mM N-acetyl-cysteine (NAC). **(D**) Same as (C), with 50 µM buthionine-sulfoximine (BSO). **(E**) time-lapse, single-cell microscopic analysis of cytoplasmic glutathione redox potential before cell death, assessed as 405/488 nm fluorescence ratio from the Grx1-roGFP2 biosensor in FL^MycER^ cells treated with OHT and IACS-1010759 (36h at the onset of filming). The time of death (t = 0) was scored based on the cellular incorporation of PI, present in the culture medium. **(F**) Immunoblot on lysates from FL^MycER^ cells treated as indicated, with the addition of 10 mM NAC concomitantly with IACS-010759. * p ≤ 0.05; ** p ≤ 0.01; *** p ≤ 0.001; **** p ≤ 0.0001.

We next sought to address whether the observed imbalances in redox homeostasis might be required for the death of OHT/IACS-010759 treated FL^MycER^ cells. Supplementation of the cultures with the ROS scavenger and glutathione precursor N-acetylcysteine (NAC) reduced the killing of double-treated cells (Figure 2C); reciprocally, the inhibitor of GSH synthesis buthionine sulfoximine (BSO) enhanced killing (Figure 2D). This effect of BSO in potentiating the cytotoxic action of IACS-010759 was confirmed in two MYC-rearranged human lymphoma cell lines, DoHH2 and Ramos (derived from a double-hit and a Burkitt’s lymphoma, respectively) (Supplemental Figure 1B). Finally, time-lapse microscopy on FL^MycER^ cells expressing Grx1-roGFP2, a biosensor of cytoplasmic glutathione redox potential^27^, revealed that an abrupt fall in GSH availability (as revealed by the increase in 405/488 nm fluorescence ratio) regularly preceded death in double-treated cells (Figure 2E, Movie S1). Altogether, the above data strongly support a causative role of oxidative stress in the IACS-010759 induced killing of MYC-overexpressing cells.

We and others previously showed that the Integrated Stress Response (ISR) drives cell death in response to IACS-010759^16,28^. Since the ISR can also be activated by oxidative stress^29,30^, we tested whether NAC could counteract the action of IACS-010759 in triggering this response, as assayed by accumulation of the ISR-associated transcription factors ATF4 and CHOP. Indeed, while both proteins readily accumulated in OHT/IACS-010759 treated cells^16^, this effect was largely abrogated in the presence of NAC (Figure 2F). Finally, antibiotics that inhibit the mitochondrial ribosome and consequently suppress OxPhos activity (e. g. tigecycline and other tetracyclines) also activate the ISR^31-34^. As with IACS-010759, tigecycline-dependent induction of the ISR was quenched by NAC in FL^MycER^ cells (Supplemental Figure 1C). These results point to a widespread role for ROS in ISR activation when interfering with OxPhos.

### Glucose and the pentose phosphate pathway maintain redox homeostasis in IACS-010759 treated cells

In all cell types examined so far, including OHT-treated FL^MycER^ cells, the cytotoxic action of IACS-010759 was suppressed by excess glucose in the culture medium^15,16,35^. Among its multiple fates, glucose serves to regenerate NADPH from NADP through the action of G6pd and Pgd, two enzymes of the oxidative phase of the pentose phosphate pathway (PPP, Figure 3A). Most relevant in this context, the conversion of NADPH to NADP serves to recycle GSSG to GSH^36^. Indeed, in parallel with the decline in GSH/GSSG ratio (Figure 2A), IACS-010759 elicited significant drops in the NADPH/NADP ratio (Figure 3B), an effect reinforced by co-treatment with OHT. Thus, we hypothesized that glucose might prevent IACS-010759 induced killing through the maintenance of redox homeostasis. In line with this concept, monitoring of the Grx1-roGFP2 biosensor revealed that the fall in GSH availability, which precedes cell death in OHT/IACS-010759 double-treated FL^MycER^ cells (Figure 2E), was reversed by adding glucose to the medium (Figure 3C).

**Figure 3.**
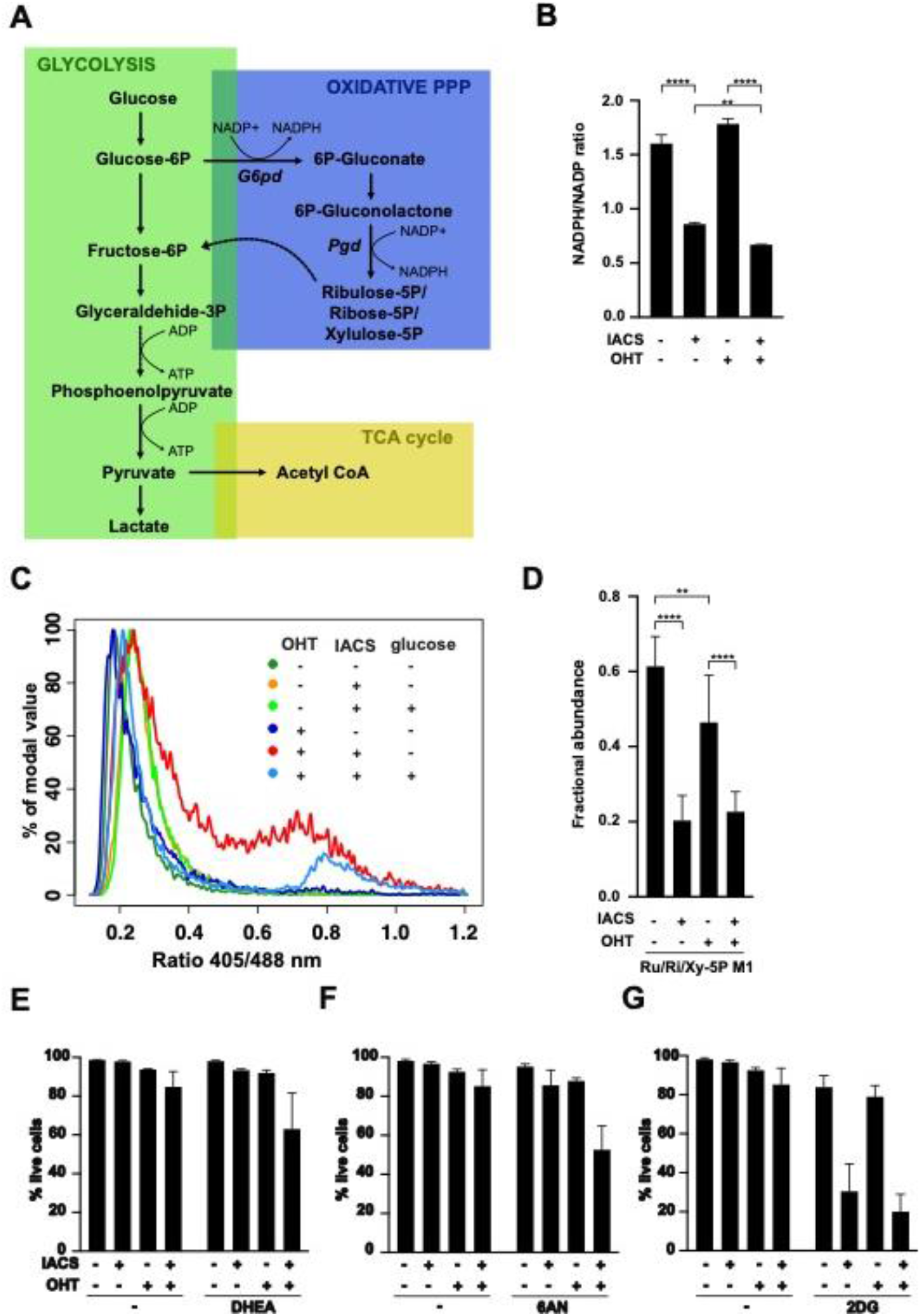
The pentose phosphate pathway protects MYC-overexpressing cells from IACS-010759-induced toxicity. (**A**) Schematic representation of glucose metabolism, highlighting NADPH- and ATP-producing reactions. (**B-G**) FL^MycER^ cells were primed with 100 nM OHT and treated with 135 nM IACS-010759, as indicated. (**B**) NADP/NADPH redox state in FL^MycER^ cells, assayed after 48 hours of IACS-010759 treatment. (**C**) cytoplasmic glutathione redox potential, assessed after 44 hours of IACS-010759 treatment in FL^MycER^ cells expressing the Grx1-roGFP2 biosensor. Where indicated, 2.75 mM glucose was added to the medium and the measurement repeated after 3 hours. (**D**) Fractional abundance of the ^13^C-labeled glucose metabolites ribulose-5P, ribose-5P and xylulose-5P (Ru/Ri/Xy-5P), the endpoints of the PPP oxidative phase, after 24 hours of IACS-010759 treatment. ** p ≤ 0.01; **** p ≤ 0.0001. (**E-G**) Cell viability after 40 hours of IACS-010759 treatment in the presence of (**E**) 50 µM dehydroepiandrosterone (DHEA), (**F**) 10 µM 6-aminonicotinamide (6AN) or (**G**) 1 mM 2-deoxyglucose (2DG). See Supplemental Figure 2E for detailed statistical analysis. Error bars: SD (n=6).

To better understand the effects of Myc and IACS-010759 on the oxidative PPP, we monitored the levels and activities of its two key enzymes, G6pd and Pgd (Figure 3A). G6pd was induced upon OHT treatment, with no effect of IACS-010759 (Supplemental Figure 2A). Yet, while OHT had only a marginal effect on G6pd enzymatic activity, IACS-010759 strongly suppressed it (Supplemental Figure 2B). Pgd instead showed no significant variations with either OHT or IACS-010759 (Supplemental Figure 2A, B). Since G6pd catalyzes the rate limiting step of the PPP, we surmised that suppression of its activity by IACS-010759 may reduce the overall glucose flux through this pathway: indeed, metabolic flux analysis with the isotopic tracer [1,2-^13^C]glucose revealed that, regardless of OHT treatment, IACS-010759 suppressed production of the final products on the oxidative PPP reactions, ribulose-5-P, ribose-5-P and xylulose-5-P (Ru/Ri/Xy-5P, Figure 3D). This decreased PPP flux would also be expected to suppress lactate M1, the isotopomer produced by glucose passing through the PPP before re-entering glycolysis (Figure 3A): while apparent in our data, this effect remained below statistical significance (Supplemental Figure 2C). On the other hand, IACS-010759 treatment increased the relative abundance of the lactate M2 isotopomer, produced by direct passage through glycolysis. Consistent with this result, the basal glycolytic rate, as assessed by glycolytic proton efflux rate (glycoPER), was potently increased by IACS-010759 (Supplemental Figure 2D). By re-routing glucose from the PPP to glycolysis, this shift would fulfill the need to maintain energetic homeostasis upon suppression of OxPhos. Of note here, while MycER activation had no measurable impact on glucose flux as determined by isotopic tracing in our experimental conditions, glycoPER analysis revealed a slight increase in basal glycolysis upon OHT treatment (Supplemental Figure 2D), in accordance with the described pro-glycolytic action of MYC^1^.

The importance of the oxidative PPP for the selective action of IACS-010759 on MYC-overexpressing cells was underscored by pharmacological inhibition of G6pd and Pgd with dehydroepiandrosterone (DHEA) and 6-aminonicotinamide (6AN), respectively^37,38^, either of these compounds enhancing cell death selectively in OHT/IACS-010759 double-treated FL^MycER^ cells (Figure 3E and Supplemental Figure 2E). In contrast, inhibiting the first step of glucose metabolism with 2-deoxyglucose (2DG)^39^ sensitized FL^MycER^ cells to IACS-010759 irrespective of OHT treatment (Supplemental Figure 2E), most likely because of the general energy impairment caused by simultaneous inhibition of OxPhos and glycolysis^40^. Finally, either DHEA or 6AN also potentiated killing by IACS-010759 in human MYC-rearranged lymphoma cells lines (Supplemental Figure 2F).

Altogether, the above data show that the NAPDH produced from glucose through the oxidative phase of the PPP (Figure 3A) protects MYC-overexpressing cells from IACS-010759 induced killing, supporting cell survival through the regeneration of GSH required to maintain redox homeostasis.

### Ascorbate potentiates the pro-oxidant and anti-tumoral effects of IACS-010759

Given the protective role of antioxidant defenses, we hypothesized that pro-oxidant agents might increase the cytotoxic action of IACS-010759. Parenteral administration of high dose of vitamin C (ascorbate) has been shown to have pro-oxidant and anti-cancer activity in preclinical models^41-44^. These results prompted several clinical trials using high dose ascorbate to treat advanced human cancer, in which ascorbate showed no activity if given as monotherapy^45,46^, but had some efficacy when combined with chemotherapy^47,48^. Importantly, all of these studies concurred to show that ascorbate had minimal toxicity. We thus tested ascorbate in combination with IACS in FL^MycER^ cells. While ascorbate had no cytotoxic effect on its own (neither with-nor without OHT priming), it significantly potentiated killing by IACS-010759, both with and without OHT (Figure 4A). Monitoring with the Grx1-roGFP2 reporter showed that ascorbate caused a progressive and profound fall of glutathione redox potential in IACS-010759 treated cells (Figure 4B, left), while those treated with ascorbate alone showed a moderate and transient reduction (2h), followed by full recovery (5h; Figure 4B, right). Comparable results were obtained in the presence of OHT (Supplemental Figure 3A).

**Figure 4.**
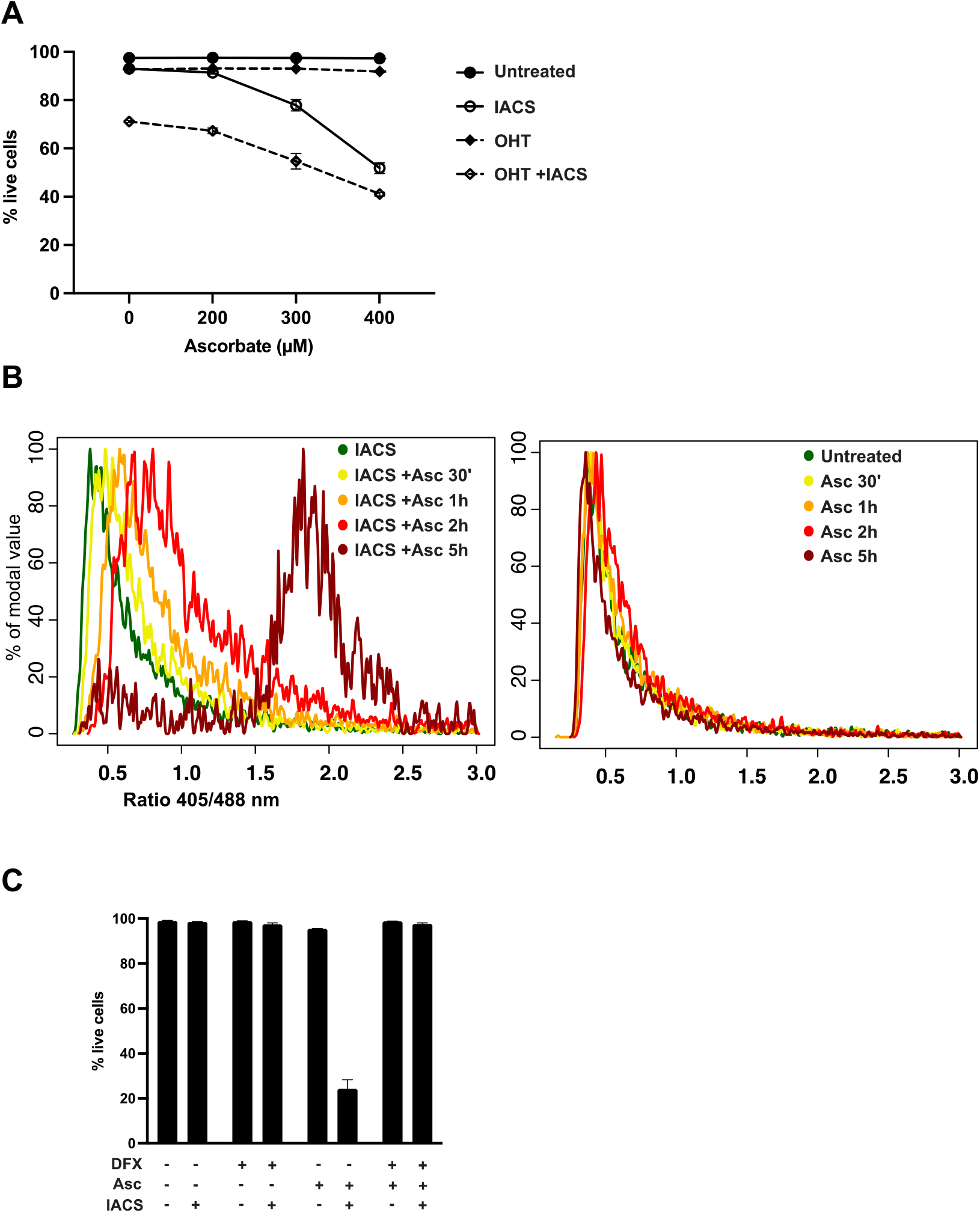
Ascorbate potentiates IACS-010759-induced cell death by increasing oxidative stress. FL^MycER^ cells were primed with 100 nM OHT (A, B) or left unprimed (C) and treated with 135 nM IACS-010759 for 36 or 42 hours, followed by addition of ascorbate (Asc, 400 µM). (**A**) Cell viability at the end of treatment (48h IACS-010759, 12h Asc). Error bars: SD (n=3). (**B**) 405/488 nm fluorescence ratio from the cytoplasmic Grx1-roGFP2 reporter in cells treated with ascorbate for the indicated periods of time, either with IACS-010759 (36h, left) or without it (right). The same experiment with OHT-primed cells is shown in Supplemental Figure 3A. (**C**) Cell viability at the end of treatment (48h IACS-010759, 6h Asc), in the presence or absence of 50 µM deferoxamine (DFX, added 1 hour before Asc). Error bars: SD (n=3).

Previous work linked the toxicity of ascorbate to ROS production through a Fenton reaction with the cell’s free iron pool^49^. Since inhibition of mitochondrial complex I can increase free iron^50^, this might explain the combined cytotoxic action of IACS-010759 and ascorbate: in line with this scenario, the levels of two mRNAs known to be destabilized by free iron, *Tfrc* and *Slc11a2*^*51*^, were reduced upon IACS-010759 treatment (Supplemental Figure 3B). Furthermore, the ferric iron chelator deferoxamine (DFX) fully prevented cell death in double-treated cells (Figure 4C). Altogether, we surmise that ascorbate sensitizes cells to IACS-010759 by reacting with labile iron, thus potentiating oxidative stress.

In line with the data obtained in in FL^MycER^ cells, ascorbate also increased IACS-010759 mediated killing in human lymphoma cell lines (Figure 5A), with the two drugs displaying significant synergistic effects (Figure 5B). To address whether these activities could be exploited in a pre-clinical setting, CD1 nude mice were transplanted subcutaneously with either Ramos or DoHH2 cells: after tumor engraftment the mice were treated with ascorbate and/or IACS-010759 over two weeks. Remarkably, the growth of either Ramos or DoHH2 xenografts was significantly delayed by the combination, but not by each drug alone (Figure 6A, B).

**Figure 5.**
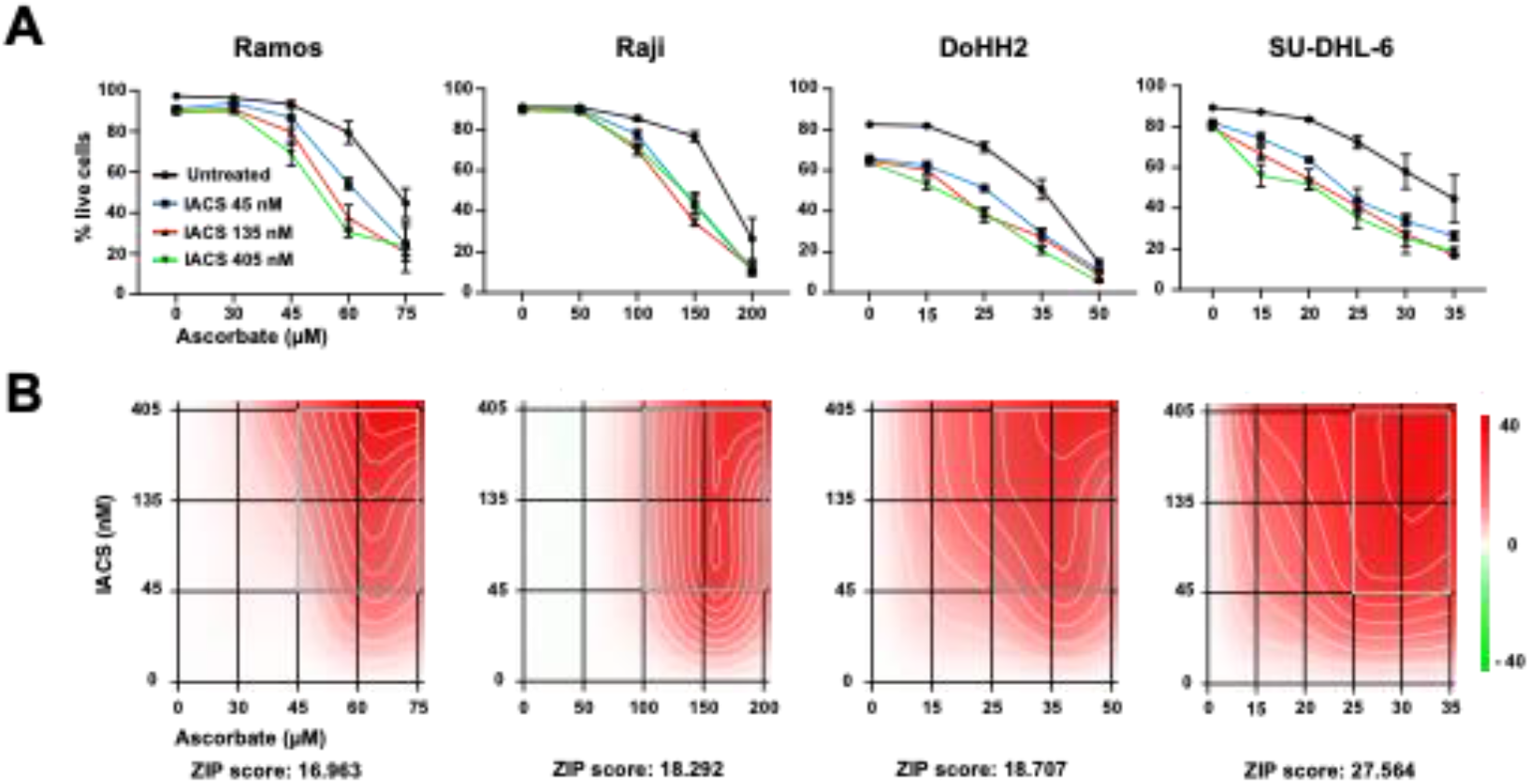
Ascorbate and IACS-010759 synergistically induce cell death of Myc-overexpressing human lymphoma cells in vitro. The indicated Burkitt’s (Ramos, Raji) and double-hit lymphoma cell lines (DoHH2, SU-DHL-6) were treated with the indicated concentrations of IACS-010759 and ascorbate, and cell viability determined after 24 hours. (**A**) Live cell counts. Error bars: SD (n=3). (**B**) Drug interaction landscapes and synergy scores, calculated according to the ZIP model: a positive ZIP score (>10) signifies a synergistic interaction. The landscape identifies the doses at which the drugs either synergize (red) or antagonize each-other (green) - the latter not observed here.

**Figure 6.**
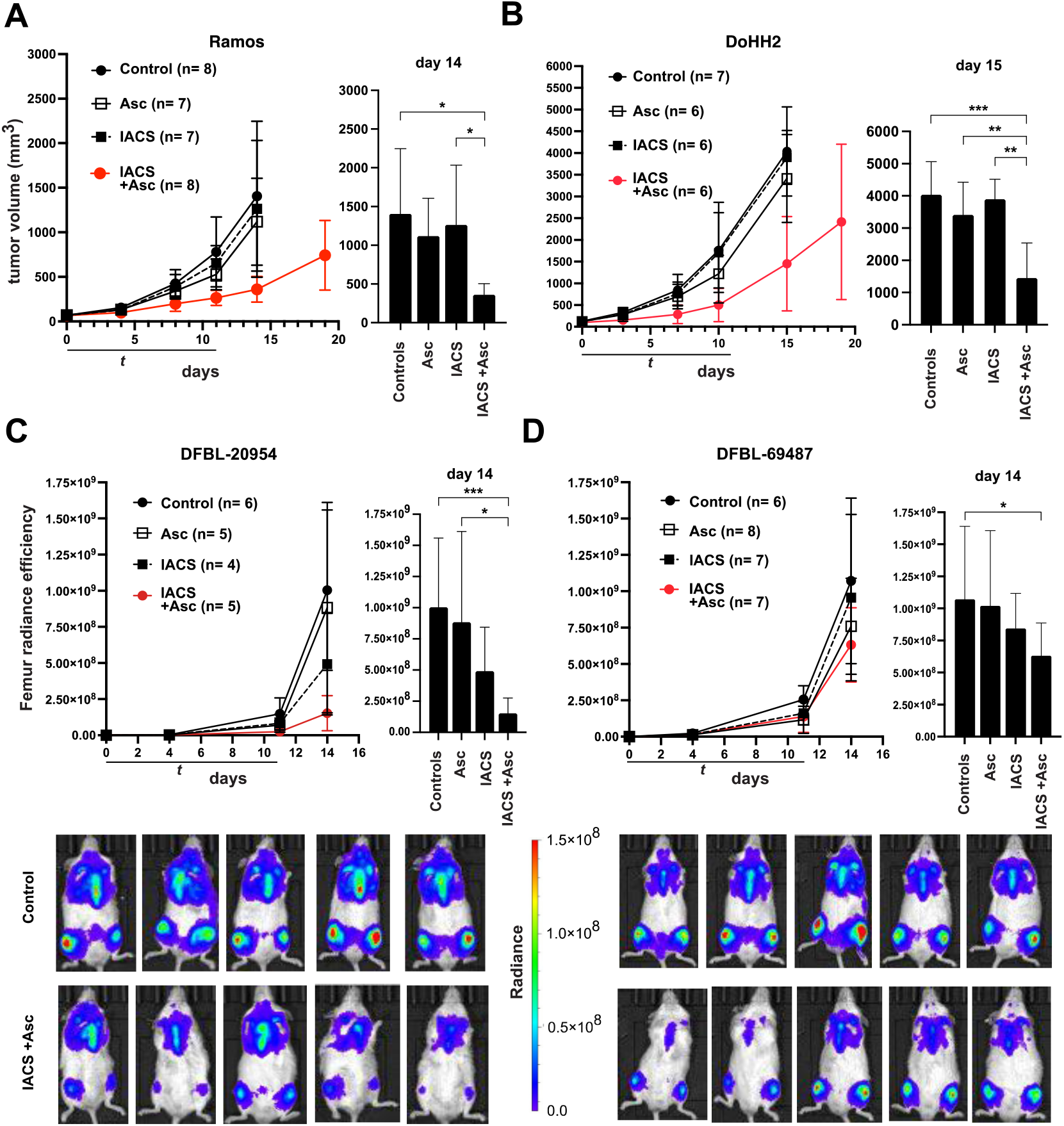
Combinatorial action of IACS-010759 and ascorbate against Myc-overexpressing lymphoma xenografts. The indicated lymphoma cell lines and PDX samples were xenografted in recipient animals and tumors allowed to develop to detectable sizes prior to randomization and treatment with IACS-010759 (IACS) by oral gavage and/or ascorbate (Asc) by intraperitoneal injection, as indicated (see Materials and Methods). Daily doses and the number of animals per group (n) are indicated in parenthesis. (**A, B**) Tumor progression (mm^3^) in CD1 nude mice bearing subcutaneous Ramos (**A**) or DoHH2 tumors (**B**). (**C, D**) Tumor progression in NSG mice injected with the luciferase-positive DHL-derived PDX lines DFBL-20954 (**C**) or DFBL-69487 (**D**), as determined by *in vivo* imaging and quantification of bilateral femur radiant efficiency. Examples form the control and double-treated groups are shown at the bottom. In each panel, the plot on the right shows the data for individual groups on the indicated day. Error bars: SD; n = animals per group. Right: one-way between-group ANOVA. * p ≤ 0.05;

We then measured the efficacy of the same drug combination against two DHL-derived patient-derived xenografts (PDX), PDX-20954 and -69487^52^, injected systemically in NSG mice. Monitoring by whole-body bioluminescence showed that growth of PDX-20954 was significantly delayed by the combination, as compared to either the control or the ascorbate-treated groups, while the effect of IACS-010759 alone did not reach statistical significance (Figure 6C). The second lymphoma, PDX-69487, showed a remarkable resistance to IACS-010759 alone even if used at a higher dose, and no significant response to ascorbate alone; nonetheless, the combination did cause a moderate, but significant reduction in tumor growth (Figure 6D).

Collectively, the above results demonstrate a potentiation of the anti-tumoral activity of IACS-010759 by ascorbate on Myc-overexpressing B cell lymphoma, identifying redox homeostasis as a valid therapeutic target to treat this type of cancer.

## DISCUSSION

One of the early transcriptome studies in DLBCL^53^ identified a disease subgroup termed OxPhos, characterized by elevated expression of genes associated with respiratory activity; further work confirmed that this classification represents a *bona fide* metabolic signature and is associated with increased levels of the antioxidant glutathione, allowing tumor cells to cope with oxidative stress^54^. We recently reported that OxPhos- and *MYC*-associated gene signatures are highly correlated in DLBCL^16^, and that over-expressed MYC sensitizes cells to inhibition of OxPhos activity, either indirectly with tigecycline^11^ or directly with IACS-010759^16^.

In the present work, we show that the MYC-mediated sensitization to IACS-010759 is brought about by a critical accumulation of oxidative stress. Ectopic MycER activation and IACS-010759 treatment in the B-lymphoid cell line FL5.12 drove the production of at least two distinct oxidative species, namely H_2_O_2_ and O_2_^•-^, which together caused a disruption of redox homeostasis and participated in activating the integrated stress response (ISR), which ultimately instigates cell death under these conditions^16^. These effects were consistently suppressed and reinforced, respectively, by the addition of compounds that up- and down-modulated glutathione synthesis, pinpointing redox unbalance as the cause of selective killing by IACS-010759.

The ability of cells to regenerate GSH from oxidized glutathione (GSSG) is dependent upon the availability of NADPH, which in turn is regenerated from NADP by utilizing different substrates: glucose in particular is the main source of NADPH in cancer cells when metabolized through the pentose phosphate pathway (PPP), under the control of G6pd and Pgd (Figure 3A)^55^. Most relevant here, the cytotoxic action of IACS-010759 in different cell types was suppressed by excess glucose in the medium^15,16,35^ and, as shown in our FL^MycER^ cells, the sensitizing effect of glucose deprivation was not always linked to an impairment in energy homeostasis, pointing to additional protective effects of this sugar^16^.

The data reported here reveal that the protective effect of glucose in MYC-overexpressing cells is mediated by its antioxidant action through the PPP pathway. First, addition of glucose reversed the loss of reduced glutathione elicited by MycER activation and IACS-010759 treatment in FL^MycER^ cells. Second, IACS-010759 treated cells showed a significant suppression of G6pd activity, as well as decreased production of Ru/Ri/Xy-5P, the final product of the oxidative phase of the PPP. Third, pharmacological inhibition of either G6pd or Pgd further enhanced the selective killing of MYC-overexpressing cells by IACS-010759. This result is in line with the previous finding that depletion of Pgd sensitized H1975 lung cancer cells to IACS-010759^17^: of note, H1975 cells bear an amplified *MYC* locus and significantly over-express the protein^56,57^. We surmise that the combined effects of MYC activity and IACS-010759 render the cells dependent upon glucose and the PPP pathway to prevent the accumulation of lethal oxidative stress.

Altogether, the aforementioned findings are consistent with the long-known pro-oxidant effects of MYC^19,20^ and reveal that these can sensitize cells to complementary pro-oxidative cues – as achieved here by OxPhos inhibition – leading to critical disruptions of redox balance. This concept led us to test whether exacerbating the disruption of redox homeostasis caused by IACS-010759 could reinforce its anti-cancer activity against MYC-driven B-cell lymphomas. Indeed, interfering with glutathione synthesis with BSO increased IACS-010759 activity on lymphoma cells. We further developed this concept by combining IACS-010759 with ascorbate (vitamin C), which has a pro-oxidant activity when injected at pharmacological doses and has shown toxicity toward diverse tumor cells, including lymphoma^42-44^. Remarkably, IACS-010759 and ascorbate synergized in killing BL and DHL cell lines *in vitro*, as well as in suppressing the growth of lymphoma xenografts *in vivo*.

The combinatorial action of IACS-010759 and ascorbate unraveled here might prove to be relevant in diverse clinical settings. First, ascorbate might increase the therapeutic window of OxPhos inhibitors, allowing their administration at clinically safe doses: this is a critical priority indeed, since a recent phase I trial revealed mechanism-related toxicity of IACS-010759, limiting its usage at effective anti-tumoral doses^58^. Similar problems were encountered with the complex I inhibitors BAY-87-2243 and ASP4132^59,60^. Second, our findings might provide a much-needed alternative treatment for some of the most aggressive forms of DLBCL that are refractory to front-line R-CHOP immune-chemotherapy. Indeed, substantial efforts are being invested toward the definition of the specific molecular features – such as oncogene translocations, mutational and transcriptional profiles – that may allow to identify those patients that will respond poorly to standard therapy, and may instead benefit from alternative regimens^7,61-65^. Among these features, MYC translocation and/or over-expression have been associated with reduced survival, especially if associated with alterations of the BCL2 oncogene^4,5^: our present results point to MYC as a biomarker for a therapeutic approach based on IACS-010759 and ascorbate.

Of note here, our previous studies highlighted therapeutic synergies between either IACS-010759^16^ or tigecycline^13^ and the BCL2 inhibitor venetoclax against the aggressive DHL subtype, characterized by joint translocation of *MYC* and *BCL2*. Having shown here that ascorbate reinforces the effect of IACS-010759 on the same DHL lines, it can be hypothesized that a triple-combination of IACS-010759, ascorbate and venetoclax may further improve the therapeutic window and treatment efficacy on these very aggressive lymphomas.

A recent report linked activation of the MYC paralogue MYCN in neuroblastoma to dependence on cysteine import and synthesis, needed to produce glutathione and avoid ferroptotic cell death^66^. Cell death induced by IACS-010759 in OHT-treated FL^MycER^ shows features typical of canonical apoptosis, like Bax/Bak-dependence and nuclear fragmentation^16^, making ferroptosis an unlikely mechanism in this setting^67^. However, a possible involvement of ferroptosis in the iron-dependent potentiation of cell killing by ascorbate remains to be investigated and – if verified – may point to a therapeutic potential of combining IACS-010759 with ferroptosis inducers.

Finally, beside DLBCL, other tumor types showed an association of MYC and/or OxPhos gene signatures with increased resistance to various therapies, including ibrutinib in mantle cell lymphoma, neoadjuvant therapy in triple negative breast cancer, and venetoclax in acute myeloid leukemia^68-70^. In all of these cases, the treatment-refractory cancer cells were shown to be sensitive to pharmacological inhibition of OxPhos: our results suggest that additional combination with a pro-oxidant drug like ascorbate may further improve the effectiveness of this therapeutic strategy.

## METHODS

### Chemicals

IACS-010759 (from the Institute for Applied Cancer Science at MD Anderson)^15^ and 4-hydroxytamoxifen (Merck Life Science, Darmstadt, Germany) were dissolved in DMSO and ethanol, respectively; sodium L-ascorbate, N-acetyl-L-cysteine (NAC), L-buthionine-sulfoximine, 6-aminonicotinamide, deferoxamine mesylate (Merck Life Science), dehydroepiandrosterone (Cayman Chemical Co., Ann Arbor, MI, USA) and tigecycline (Carbosynth, Newbury, UK) were dissolved in water.

### Cell lines

The human lymphoma cell lines were DoHH-2 (CVCL_1179), SU-DHL-6 (CVCL_2206), Ramos (CVCL_0597) and Raji (CVCL_0511), and the murine B-lymphoid cell line FL5.12 (CVCL_0262) expressing MycER™ (FL^MycER^)^16^, were grown in regular RPMI medium (Euroclone, Pero, Italy), which includes 2mM glutamine and 11 mM glucose, supplemented with 10% fetal bovine serum (FBS) and, for FL5.12 cells only, 2 ng/mL murine interleukin 3 (PeproTech, Rocky Hill, NJ, USA). Prior to experiments involving IACS-010759, all cells were passaged in glucose-free RPMI-1640 medium (Thermo Fisher Scientific, Waltham, MA, USA), which includes 2mM glutamine, supplemented with 10% FBS and 2.75 mM glucose. To induce MycER activity, 100 nM 4-hydroxy-tamoxifen (OHT; Merck) was added to the medium 48 hours before treatment with IACS-010759 or other drugs, as indicated. All cells were incubated at 37° C in a humidified air atmosphere supplemented with 5% CO2. SU-DHL-6, Raji and Ramos cell lines were imported from the ATCC repository (https://www.lgcstandards-atcc.org); DoHH-2 cells were imported from the DSMZ repository (https://www.dsmz.de); FL5.12 cells were a gift from Pier Giuseppe Pelicci. All lines were stocked and made available by IEO’s core Tissue Culture facility, where they were also validated and tested for mycoplasma infection.

### Biochemical assays

GSH/GSSG and NADPH/NADP were quantified with luminescence-based assay kits (Promega, Madison, WI, USA), and the enzymatic activities of G6pd and Pgd with colorimetric assay kits (BioVision/Abcam, Waltham, MA, USA), according to manufacturer’s specifications.

### Time-lapse analysis of the Grx1-roGFP2 biosensor

Retroviral pLPCX vectors expressing the mitochondrial or cytoplasmic form of Grx1-roGFP2^27^ and roGFP2-ORP1^23^ were obtained from Addgene (Addgene, Watertwon, CA, USA) and used to transduce FL5.12^MycER^ cells, followed by a one-week selection with puromycin (AdipoGen AG, Liestal, CH) 1.5 µg/mL. Cells expressing either form of Grx1-roGFP2 were primed with OHT and treated with IACS-010759 for approximately 36h, and then transferred to glass bottom petri dishes (MatTek Corporation, Ashland, MA, USA) in the continuous presence of the drugs, with addition of 1 µg/mL PI to mark dead cells. The cultures, kept at 37 °C and 5% CO2, were imaged with a Leica SP8 FSU confocal microscope (Leica Microsystems, Wetzlar, Germany) through a 63x/1.4NA oil immersion objective lens. The oxidized and reduced forms of the Grx1-roGFP2 biosensor were excited by the 405 nm and 488 nm laser lines, respectively. The emitted fluorescence for both forms was collected with an opened pinhole (2.7AU) in the 500-540 nm acquisition window and sequentially acquired with the same HyD detector set in counting mode with a pixel size of 180 nm. The PI molecule was excited with the 561 nm laser line and the emitted signal collected between 590 and 700 nm. Images were acquired every 5 minutes for 12 hours.

The ratio between the emission intensities of oxidized and reduced Grx1-roGFP2 was calculated thanks to a custom-made ImageJ/Fiji macro. Briefly, after background subtraction, the combination of the 2 signals was obtained and the resulting image (combined image) was filtered with a Gaussian filter and segmented with the Otsu algorithm. The resulting mask was applied to the original images of oxidized and reduced Grx1-roGFP2 to obtain the ratio image. To segment the single cells in the images at fixed time-points, the combined image was segmented using a more permissive algorithm (Li) and the resulting binary image subjected to the “analyse particle” function of ImageJ. The region of interest (ROI) values of single cells were then used to calculate the oxidized/reduced Grx1-roGFP2 fluorescence ratio in every segmented cell. Single cells were manually tracked to record the Grx1-roGFP2 ratio and PI signal intensity over time.

### Flow Cytometry

Flow cytometry was conducted on a MACSQuant Analyzer (Milteniy Biotec, Bergisch Gladbach, Germany) and data analyzed with FlowJo software (version 10.6.1; BD Biosciences, Franklin Lakes, NK, USA). For cell viability counts, cells were resuspended in ice-cold PBS in the presence of Propidium iodide (PI) 1 µg/mL. Viability was expressed as percentage of live (PI negative) cells on total cells counted.

To quantify the fluorescence from Grx1-roGFP2 and roGFP2-ORP1 biosensors, the oxidized and reduced forms were analyzed with the V2 (405/585-40 nm) and B1 (488/525-50 nm) channels, respectively. Immediately before counting, PI 1 µg/mL was added to the medium to exclude dead cells.

Quantification of superoxide anion was performed by staining live cells with 10 µM Dihydroethidium (Thermo Fisher Scientific): following 30 minutes of incubation with the reagent, cells were washed and resuspended in Phosphate-buffered saline for flow-cytometric analysis. Immediately before counting, 4′,6-diamidino-2-phenylindole (DAPI; Merck Life Science) 0.1 µg/mL was added to the medium to exclude dead cells.

### Immunoblot analysis and cell fractionation

Immunoblot analysis was performed as previously described^16^. For the analysis of subcellular fractions, the same numbers of cells from each sample were lysed 5’ on ice with NP-40 fractionation buffer (10 mM Hepes pH 7.4 1, 250 mM Sucrose, 25 mM KCl, 2 mM MgCl2, 1 mM EGTA, 0.1% NP-40, 1 mM PMSF). This total lysate was then centrifuged at 1000x g for 5 minutes, and the supernatant collected as the cytoplasmic fraction. The nuclear fraction was obtained by washing the pellet twice with fractionation buffer without NP-40 and finally resuspending it in Laemmli buffer. The following antibodies were used: mouse monoclonals against vinculin (Merck Life Science) and CHOP (Cell Signaling Technology, Danvers, MA, USA); rabbit monoclonals against ATF4, Nrf2, GAPDH, Histone H3, G6pd (Cell Signaling Technology) and Pgd (Abcam, Cambridge, UK).

### Basal glycolysis rate measurement

Basal glycolysis was assessed on a Seahorse XFe96 Analyzer (Agilent Technologies, Santa Clara, CA, USA) using Agilent Seahorse XF Glycolytic Rate Assay kit, following the manufacturer’s instructions. Before the assay, FL^MycER^ cells were treated for OHT and IACS-010759 for 72 and 24 hours as indicated. The cells were then counted and attached to 96-well Seahorse cell culture microplates, pre-coated with Corning™ Cell-Tak (Life Sciences) according to manufacturer’s instructions, at a density of 80,000 cells per well, in XF RPMI Medium pH 7.4 with 1 mM HEPES (Agilent Technologies) supplemented with 2.75 mM glucose, 1 mM sodium pyruvate, and 2 mM L-glutamine. The plate was incubated at 37 °C for 1 h in a non-CO2 incubator. After ECAR baseline measurements, 0.5 μM rotenone plus 0.5 μM antimycin A and 50 mM 2-deoxy-glucose (2-DG) were added sequentially to each well. The results were analyzed using the Seahorse Wave Desktop Software Version 2.6 (Agilent Technologies) and normalized by cell number using CyQUANT Cell Proliferation Assay (Thermo Fisher Scientific). Data were exported into the XF Report Generator for calculation of the parameters from the Glycolytic Rate Assay. Results are mean ±SD of minimum 8 technical replicates, and representative of 2 independent experiments.

### Metabolic flux analysis

For metabolic flux analysis, cells were exposed for 4 hours to 1.5 mM D-[1,2-^13^C2]glucose in complete medium containing 1.5 mM glucose. Cells were then harvested, washed twice in ice-cold PBS and the pellets snap frozen in liquid nitrogen. Pellets were then resuspended in 250µl methanol/acetonitrile 1:1 and spun at 20,000g for 5 min at 4°C. Supernatant were then passed through a regenerated cellulose syringe filter, dried and resuspended in 100µl of MeOH for subsequent analysis.

We used an ExionLC™ AC System (AB Sciex, Framingham, MA, USA) coupled with an API-3500 triple quadrupole mass spectrometer (AB Sciex). Quantification of different metabolites was performed using a cyano-phase LUNA column (50mm x 4.6mm, 5 µm; Phenomenex, Torrance, CA, USA) by a 5 min run in negative ion mode. Mobile phases were: (A) water and (B) 2 mM ammonium acetate in MeOH; the gradient was isocratic 90% B with a flow rate of 500 µl/min. MultiQuant™ software (version 3.0.2, AB Sciex) was used for data analysis and peak review of chromatograms. The identity of all metabolites was confirmed using pure standards.

### RNA-seq analysis

The RNA-seq data and Differentially expressed genes (DEG) called in cells treated with OHT and/or IACS-010759 were described in our previous work^16^ and are accessible through NCBI’s Gene Expression Omnibus (GEO; RRID: SCR_005012; GSE149073). DEG Canonical Pathway Analysis to identify the most likely activated pathways (based on z-score) was performed with the Ingenuity Pathway Analysis software package (QIAGEN, Venlo, Netherlands; RRID: SCR_008653).

### Xenograft models and treatment

5×10^6^ DoHH-2 or Ramos cells were xenografted subcutaneously in sub-lethally irradiated (3 Gray), 8 weeks old, female CD1-nude nu/nu mice (Envigo Indianapolis, IN, USA), and expanded by serial subcutaneous transplantation of tumor fragments. Tumors were allowed to grow for about 10-14 days, followed by exclusion of outliers, randomization of the experimental groups and start of treatments (days 0 to 11). Tumor volumes were assessed from the start of the treatment every two days with a digital caliper and calculated as 1/2 length×width^2^ (mm^3^). The following treatment schemes were used: daily oral gavage with IACS-010759 (1 or 2.5 mg/kg, as indicated in the figures) for 5 days, followed by two days off and a repeat of the same scheme, for a total of 12 days; intraperitoneal injection of ascorbate (4 or 8 g/kg) twice a day (separated by 8 hours) for 5 days, followed by two days off and a repeat of the same scheme. IACS-010759 was suspended in 0.5% methylcellulose (Merck Life Science), while ascorbate was dissolved in physiological saline.

The DHL patient-derived xenografts (PDX) DFBL-20954-V3-mCLP and DFBL-69487-V3-mCLP^52^ were obtained from the Dana Farber Cancer Institute Center for Patient Derived Models (CPDM). 1×10^6^ cells were xenografted via tail vein injection into 8 weeks old, male NSG mice (Charles River, Calco, Italy). Tumor engraftment was confirmed 7 days after transplant by whole body imaging on an IVIS Lumina III platform following intra-peritoneal injection of 150 mg/kg XenoLight D-Luciferin (PerkinElmer, Waltham, MA, USA) and anesthesia with isoflurane. The animals were subsequently randomized in the different experimental groups to start the treatment with IACS-010759 and/or ascorbate, as described above. The response to treatment was assessed by whole body imaging at days 4, 11 and 14. The data were analyzed with the Living Image Software, version 4.2 (Caliper Life Sciences, Hopkinton, MA, USA). Radiant efficiency was quantified bilaterally on femurs based on the epifluorescence signal as indicated in the user manual.

Experiments involving animals were done in accordance with the Italian Laws (D.lgs. 26/2014), which enforces Dir. 2010/63/EU (Directive 2010/63/EU of the European Parliament and of the Council of 22 September 2010 on the protection of animals used for scientific purposes), and authorized by the Italian Health Ministry with project nr. 173/2021-PR. Mice were housed in individually ventilated caging (IVC) systems (Sealsafe Plus, Tecniplast, Buguggiate, Italy), on autoclaved sawdust bedding (Lignocel® 3/4; Rettenmaier & Sohne, Ellwangen-Holzmühle, Germany), provided with autoclaved diet (VRF1 (P), SDS, Witham, UK) and autoclaved water *ad libitum*. Animals were handled and treated in laminar flow hoods (CS5 Evo and BS48, Tecniplast, Buguggiate, Italy).

### Quantification and statistical analysis

Drug interaction landscapes and delta scores for synergy were based on the ZIP model in SynergyFinder^71^. Comparisons between treatments for *in vitro* cell culture and biochemical experiments were carried out by one-way or two-way ANOVA. Partitioning the interaction effects in two-way ANOVA allowed us to test for particular differences among treatments. Between treatments tests for *in vivo* tumor growth were performed by one-way ANOVA. Comparisons between treatments tests for *in vivo* tumor growth were performed by one-way ANOVA. When pairwise comparisons of means were needed, post-hoc tests were made according to Tukey’s procedure. GraphPad PRISM 8 (RRID: SCR_002798) or R (RRID:SCR_001905) software were used for all analyses. The numbers of independent biological replicates are indicated in the figures.

## Acknowledgements

We thank Stefano Campaner, Oskar Fernandez-Capetillo, Gioacchino Natoli and members of the Amati lab for insightful comments and discussions, Alberto Gobbi and Manuela Capillo for their help with the management of mouse colonies, Simona Ronzoni for her help with flow cytometry, and Marzia Fumagalli and Claudia Iavarone for help with material transfer agreements. We are grateful to Christopher Vellano, Joseph Marszalek and Giulio Draetta for providing IACS-010759. This work was supported by a grant from the Italian Association for Cancer Research (AIRC, IG2018-21594) and ERA Permed (2020-090) to B. Amati, and by institutional funds from the Ricerca Corrente and 5×1000 programs of the Italian Ministry of Health. G. Donati was supported by a post-doctoral fellowship from the Fondazione Umberto Veronesi.

## Authorship contributions

GD organized and performed most of the experiments, with technical assistance from PN and AV. VV and AG contributed part of the *in vitro* experiments on cellular drug responses, MA and NM with metabolomic analysis, LC and AB with the glycoPER analysis, GB with statistical analysis, OC with computational and graphical tools, and SR with microscopy experiments and image analysis. GD and BA designed and coordinated the project, and wrote the manuscript.

---

**Supplemental Figure 1.**
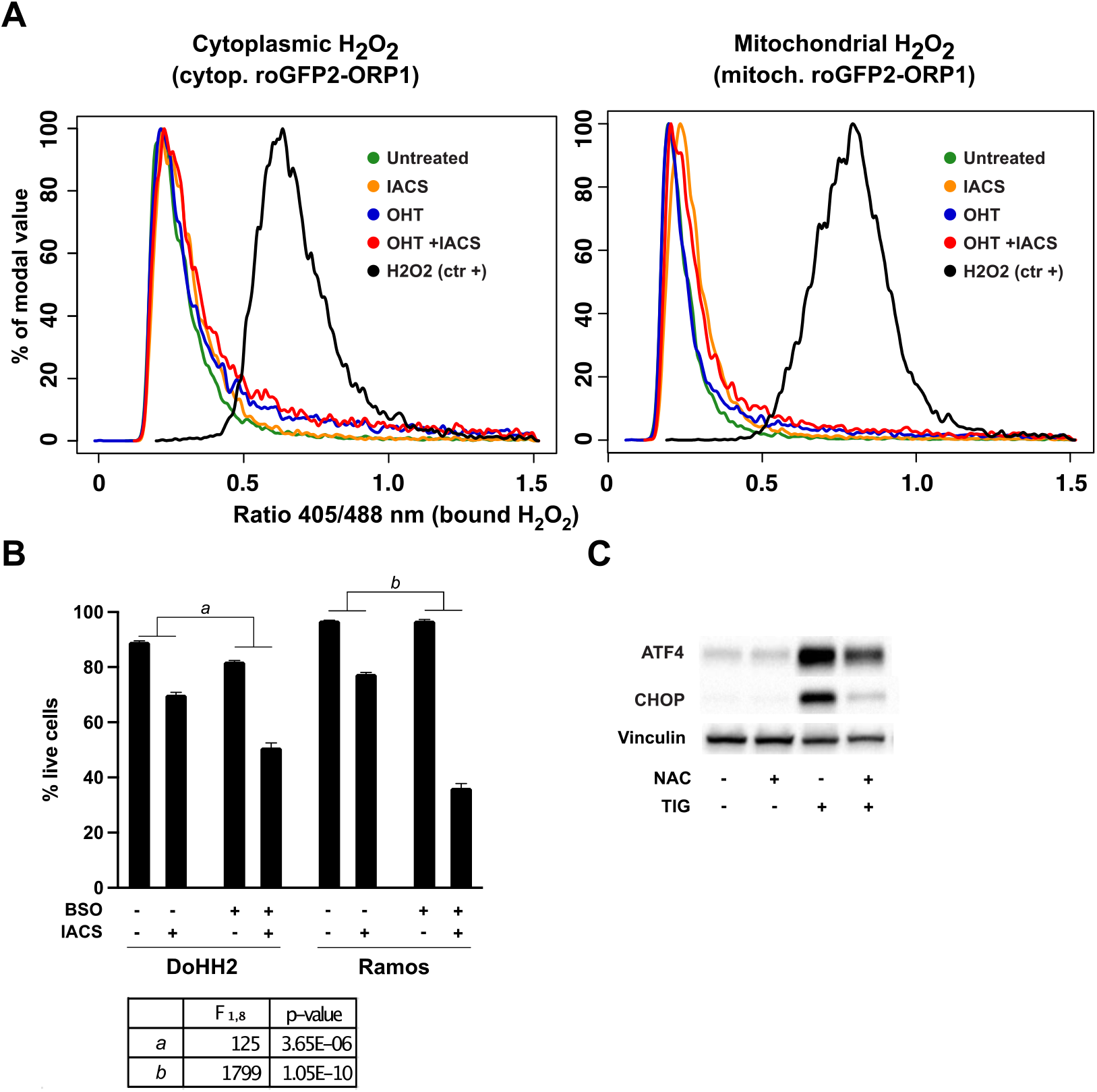
Reactive Oxygen Species contribute to the effects of MYC and mitochondrial inhibitors. (**A**) Representative distribution plots for the 405/488 nm fluorescence ratio with the cytoplasmic (left) and mitochondrial (right) roGFP2-ORP1 biosensor in OHT- and/or IACS-010759-treated FL^MycER^ cells (as in Figure 1C). As a positive control, cells were treated with 200 µM H2O2 immediately before cytometric analysis as positive control (ctr +). (**B**) Cell viability of DoHH2 and Ramos lymphoma cells treated where indicated with 135 nM IACS-010759 for 40 hours, either alone or in the presence of 50 µM BSO. Error bars: SD (n=3). The table shows the two-way ANOVA analysis for IACS-010759-induced cell death in the presence and absence of BSO. **(C**) Immunoblot on lysates from FL^MycER^ cells treated with 5 µM tigecycline (TIG) for 48 hours, in the presence or absence of 10 mM NAC, as indicated.

**Supplemental Figure 2.**
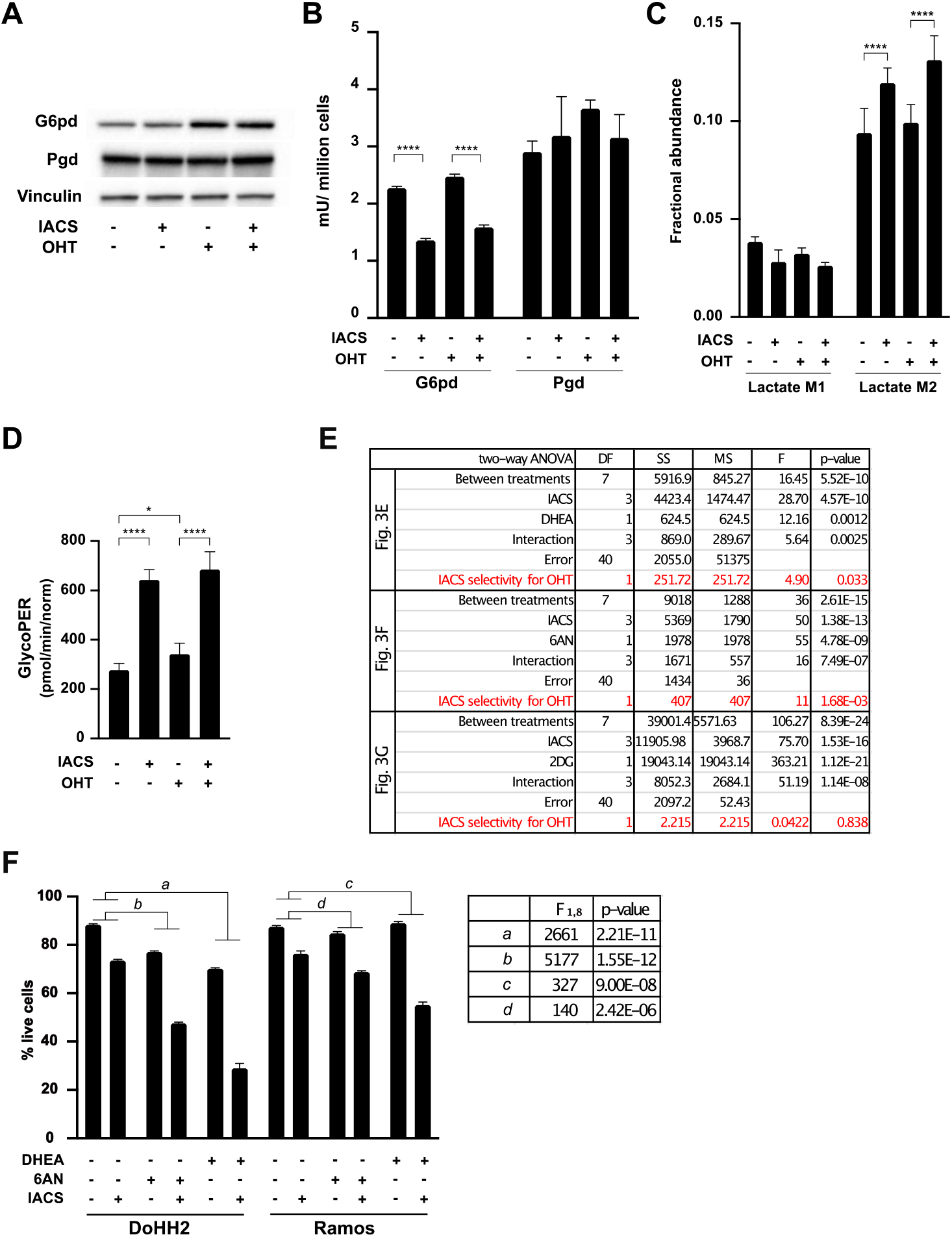
The pentose phosphate pathway protects MYC-overexpressing cells from IACS-010759-induced toxicity. FL^MycER^ cells were primed with 100 nM OHT (48h) and/or treated with 135 nM IACS-010759, as indicated. (**A**) Immunoblot analysis and (**B**) Enzymatic activity assays for G6pd and Pgd after 24 hours of IACS-010759 treatment. (**C**) Fractional abundance of ^13^C-labeled lactate, the endpoint of the anaerobic glycolytic pathway, after addition of [1,2-^13^C]glucose to FL^MycER^ cells treated as in (A). (**D**) Basal glycolytic rate, calculated form the glycolytic proton efflux rate (glycoPER), in FL^MycER^ cells treated as in (A). * p ≤ 0.05; **** p ≤ 0.0001. (**E**) Two-way ANOVA analysis of the data shown in Figure 3E, 3F and 3G, as indicated. A test for the selective cytotoxicity of IACS-010759 toward OHT-treated FL^MycER^ cells is reported in red at the end of each section for the co-treatments with DHEA, 6AN and 2DG, respectively. (**F**) Cell viability of DoHH2 and Ramos lymphoma cells treated with 135 nM IACS-010759 (40h), either alone, with 50 µM DHEA, or with 10 µM 6AN. Error bars: SD (n=3). The table shows the two-way ANOVA analysis of IACS-010759-induced cell death in the presence and absence of either DHEA or 6AN.

**Supplemental Figure 3.**
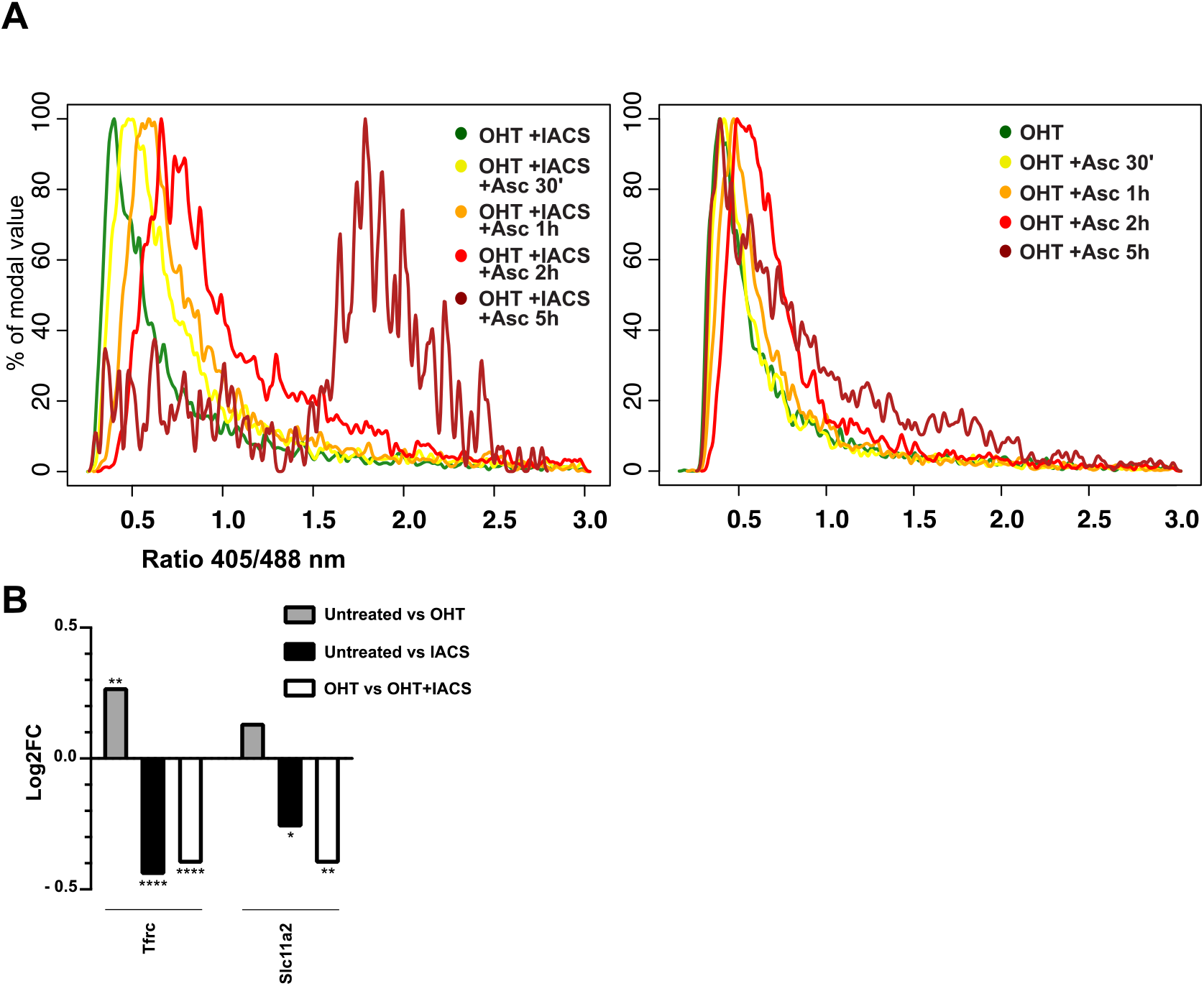
Ascorbate potentiates IACS-010759-induced cell death by increasing oxidative stress. (**A**) As Figure 4B, in OHT-primed cells. (**B**) Log fold change (Log2FC) of the iron-regulated mRNAs *Tfrc* and *Slc11a2* in FL^MycER^ cells with 135 nM IACS-010759, with or without pre-treatment with OHT, based on our previous RNA-seq profiles^16^, with the following q-values: * q ≤ 0.05; ** q ≤ 0.01; **** q ≤ 0.0001.

